# Reflexive gaze following in common marmoset monkeys

**DOI:** 10.1101/682971

**Authors:** Silvia Spadacenta, Peter W. Dicke, Peter Thier

**Affiliations:** Hertie Institute for Clinical Brain Research, Department of Cognitive Neurology, Otfried-Müller-Str. 27, 72076, Tübingen, Germany

**Keywords:** Marmoset, Face, Gaze following, Joint Attention

## Abstract

The ability to extract the direction of the other’s gaze allows us to shift our attention to an object of interest to the other and to establish joint attention. By mapping one’s own expectations, desires and intentions on the object of joint attention, humans develop a Theory of (the other’s) Mind (TOM), a functional sequence possibly disrupted in autism. Although old world monkeys probably do not possess a TOM, they follow the other’s gaze and they establish joint attention. Gaze following of both humans and old world monkeys fulfills Fodor’s criteria of a domain specific function and is orchestrated by very similar cortical architectures, strongly suggesting homology. Also new world monkeys, a primate suborder that split from the old world monkey line about 35 million years ago, have complex social structures. One member of this group, the common marmoset (Callithrix jacchus), has received increasing interest as a potential model in studies of normal and disturbed human social cognition. Marmosets are known to follow human head-gaze. However, the question is if they use gaze following to establish joint attention with conspecifics. Here we show that this is indeed the case. In a free choice task, head-restrained marmosets prefer objects gazed at by a conspecific and, moreover, they exhibit considerably shorter choice reaction times for the same objects. These findings support the assumption of an evolutionary old domain specific faculty shared within the primate order and they underline the potential value of marmosets in studies of normal and disturbed joint attention.

**HIGHLIGHTS:**

- Common marmosets follow the head gaze of conspecifics in order to establish joint attention.
- Brief exposures to head gaze are sufficient to reallocate an animal’s attention.
- The tendency to follow the other’s gaze competes with the attractional binding of the conspecific’s face

## RESULTS AND DISCUSSION

Common marmosets are well known for having a peculiar interest in faces [1,2]. Unlike macaques, the species of old world primates studied best, and other non-human primate species, they often engage in mutual gaze, for example in the context of joint action tasks [3]. Many individuals even seek eye contact with their human caretakers (personal observations). Common marmosets also care about the orientation of a human face as demonstrated by the fact that human head-gaze biases choices in an object selection task [4]. While this latter behavior may indicate an inherent capacity for gaze following, it remains to be shown that it can also be triggered by conspecifics. By the same token the lack of high resolution behavioral data has as yet precluded well-founded inferences about the relationship of marmoset gaze following to gaze following exhibited by humans and rhesus monkeys, the two species of old world primates for which detailed behavioral and neuronal data are available [5,6]. Gaze following of macaques and humans is reflex-like in the sense that it is fast and hard to suppress, two features that have contributed to the assumption of a domain specific faculty [7–12] based on a dedicated neural system [13]. Do marmosets follow the gaze of conspecifics in the same reflex-like manner? An affirmative response would support the notion that gaze following in extant primate lines is homologous, i.e. a reflection of shared ancestry.

In order to address these questions, we trained 3 common marmosets (2 females, 1 male) to execute a free choice task in a well-controlled experimental setup that allowed us to head-restrain the animals to precisely track eye movements. A conspecific’s face, oriented either to the left or to the right, was presented on a monitor for a variable time ranging between 100 and 600 ms in steps of 100 ms (see figure 1A) and the observing animal was allowed to scrutinize the face with eye movements confined by the boundaries of the portrait. The facial portrait was followed by the appearance of two targets placed at −5° and +5° from the center on the horizontal axis. The animals had to freely choose one of the two targets, a human face (2° × 3°extension), by making an indicative saccade into a window of 2° centered on the target within 500 ms. Independent of the orientation of the conspecific’s face, both possible choices were rewarded, provided that the eyes had met the fixation requirements.

**Figure 1.**
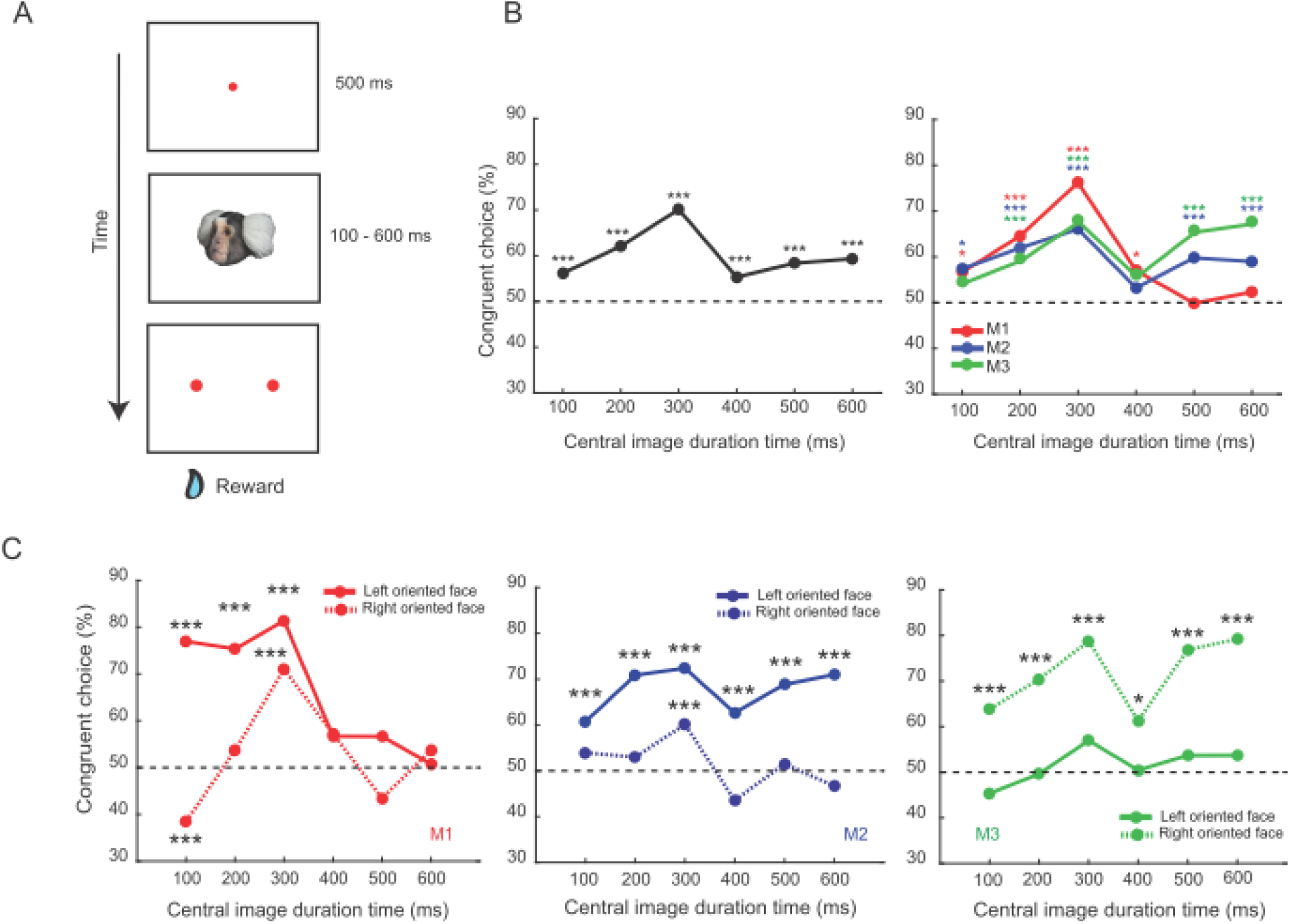
Oriented faces bias the animal’s choices to targets congruent with gaze direction. (A) Behavioral paradigm. The trial started with the presentation of a central fixation dot. Once fixation was established, the oriented face of a conspecific (replaced by other stimuli in control experiments) appeared for a variable time (100 – 600 ms in steps of 100 ms). The disappearance of the conspecific’s face or the control stimuli and the simultaneous appearance of two peripheral targets was the go signal for the animals to freely choose one of two peripheral targets presented on the horizontal axis at 5° right and left of the center respectively, by means of a saccade. The animal received a reward if the fixation requirements were met. The fixation window for the central dot had a size of 2°×2°, a size of 2°×3° for the peripheral targets portraits of a human (represented in figure as red dots) and for the central portraits/control stimuli it corresponded to the extent the centrally presented image (7° × 6°). (B) Left panel: plot of the percentage of target choices congruent with portrait orientation as function of the duration of presentation. Pooled data (monkeys M1, M2, M3). Binomial probability: *** p < 0.001 Right panel: plot of the percentage of target choices congruent with portrait orientation for the individual animals M1, M2 and M3. Binomial probability: * p < 0.05, *** p < 0.001 (C) Monkey specific plots of the percentage of target choices congruent with portrait orientation, separating portraits oriented to the left and right respectively. In each panel (left, M1; center, M2; right, M3) the solid line stands for left oriented portraits and the dashed one for right oriented portraits.

### Common marmosets follow the gaze of a conspecific in a quasi-reflexive manner

Figure 1B, left panel, plots the percentage of target choices in the direction of the face orientation (“congruent choices”) as function of the duration of the availability of the portrait. The graph depicts data pooled over all three animals and the two possible face orientations: congruent choices exceeded chance level significantly (binomial probability test, p < 0.05), indicating that the observing monkey tended to follow the gaze of the portrayed monkey. This preference was already apparent after a presentation duration of the portrait of only 100 ms and got stronger for longer presentation durations peaking at 300 ms exposure time (see also S1). This dependence on exposure duration is similar to the one exhibited by human observers when exposed to symbolic central cues such as pointing arrows. They typically demonstrate a gradual buildup of their spatial target preferences cued by central stimuli, reaching an optimum at 300 ms [14,15]. As shown in the right panel of figure 1B, the dependence of the choice bias on presentation duration was the same in all three animals for up to 300 ms. Only later, the individual plots start to diverge: interestingly, two of our animals (M2 and M3) showed a clear second peak, overall conveying the impression of an oscillatory pattern with a period of about 250 - 300 ms. Periodic fluctuations of attention between two locations with a period of 4 HZ have also been described for human and macaque spatial vision [16,17]. Yet, given the fact that the third animal exhibited a different pattern, characterized by an absence of a second gaze following peak and a constant choice at chance level after 300 ms, further studies will be needed to critically assess the possibility of periodicity.

All three animals exhibited individually different directional biases for the left and the right respectively, modifying their choice behavior on top of the influence of directional information provided by facial orientation. Directional biases became apparent when plotting the dependence of choice preference on head gaze direction for the three individual animals independently for head gaze to the left and to the right (Figure 1C). For example, a bias to the left side boosted the correct responses for the left oriented head gaze portraits (M1 and M2, left and central panel respectively), and for the right oriented when the bias fell on the other side (M3, right panel). Nonetheless, the bias never altered the overall response curve shape with a peak for congruent choices at around 300 ms. A significant dominance of congruent choices peaking at 300 ms could be seen in M1 and M2 even for congruent choices prompted by portraits oriented towards the animal’s non preferred side (binomial probability, p < 0.001). A comparable tendency in M3 did not reach significance (binomial probability, p = 0.1). The basis of the directional bias remains unclear. The fact that it differs between individuals indicates that hidden imbalances in the setup that might bind attention can hardly matter.

When the animals were confronted with direct gaze of a conspecific with the face turned straight or alternatively, with black or grey disks of a similar size, likewise lacking directional information, target choices of all three animals did not differ significantly from chance level at most exposure times with the exception of the shortest one (Figure 2A, pooled data). In particular the choice peaks for 300 and 600 ms could no longer been seen. For 100 ms exposure, overall pooled choices to the left were significantly more frequent than to the right. Two of the individual monkeys (M1, M2) exhibited this preference for the left, whereas the third one (monkey M3) a preference for the right. The individual directional preferences for the left and the right corresponded to the direction of the biases seen in the responses to oriented gaze (Figure 1C), yet, now confined to the shortest exposure only. We think that the disappearance of the directional bias for longer exposure times might be a consequence of increasing attraction towards the central object, overriding the bias, no matter if the central object is the neutral disk or the portrait of a conspecific looking straight. This interpretation has interesting implications for the experiments with oriented faces, which showed a persistence of the directional biases independent of exposure time. Here the directional gaze seems to suppress the buildup of attraction to the central object, facilitating the readiness to look elsewhere as determined by the resultant of the other’s gaze direction and an internal directional bias.

**Figure 2.**
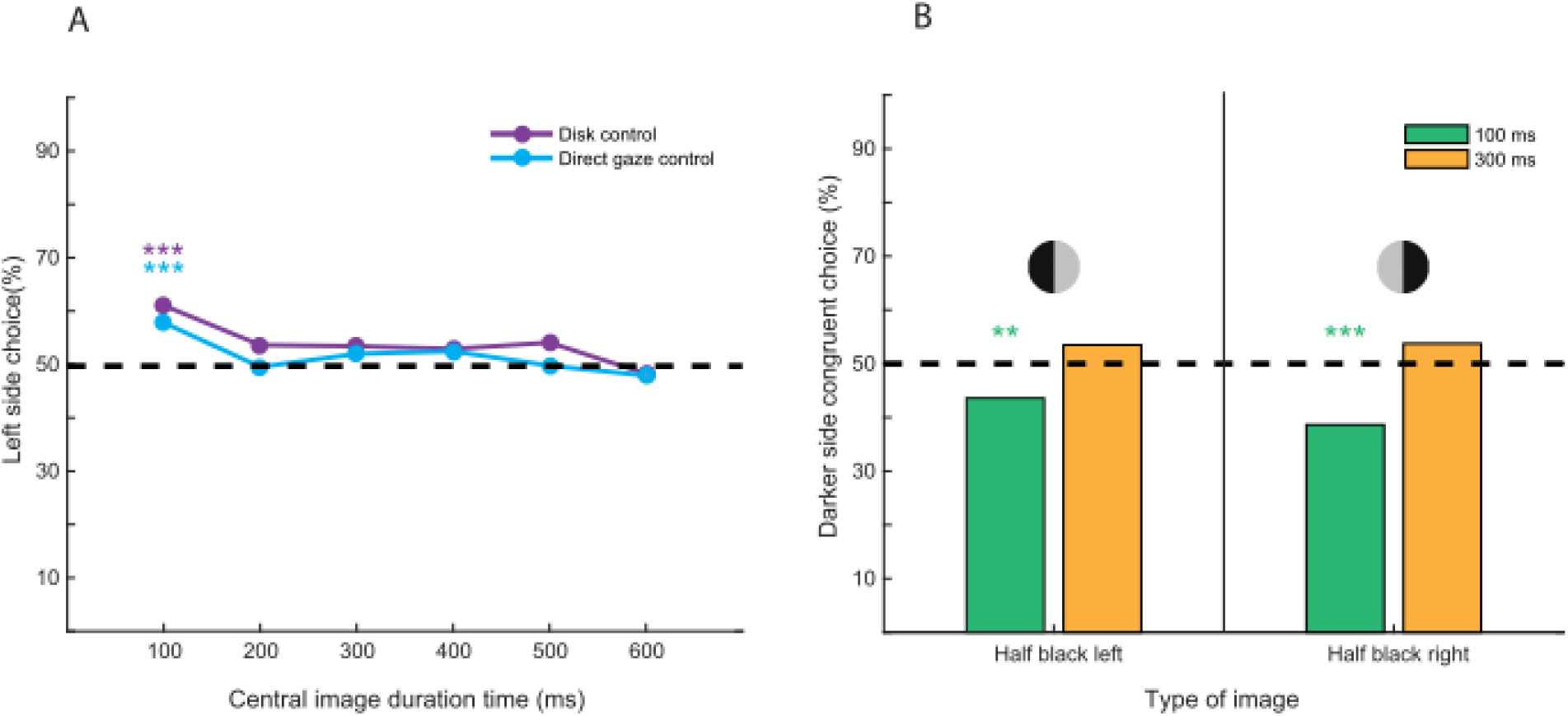
Direct gaze and disk stimuli attract the animals’ attention towards the center. (A) Plot of left target choice as function of the duration of a central stimulus, either a conspecific’s face looking straight at the observer (direct gaze) or a circular grey or black disk of similar size. Data pooled over the three animals (monkeys M1, M2, M3). A choice bias is evident at the shortest duration time, whereas for longer exposures the animals chose the targets on the right and left at random. Binomial probability: *** p < 0.001. (B) Bar plot of percentage of choices congruent with the darker half of a bipartite disk. Data pooled over the three animals (monkeys M1, M2, M3) and choice direction. For 100 ms presentation duration, the animals exhibited a significant preference for the target on the brighter side, i.e. opposite to the side preference to be expected based on a mechanism exploiting the luminance asymmetry associated with face orientation. At 300 ms the choices did not indicate a preference. Binomial probability: ** p < 0.01; *** p < 0.001.

The white ear tufts on the left and right of the darker central face of a straight ahead marmoset offer a symmetric luminance profile. Once the animal turns the head to the side, symmetry is lost as the visible area of the ear tuft on the side of the head turn will decrease, whereas the area of the other one will increase (see figure S3). Hence, gauging the extent of the luminance asymmetry may be a simple way to determine the other’s head gaze direction without the need to process other aspects of the face. To test whether left-right differences in the luminance of an object prompt an orienting response of the observer, we exposed all 3 animals to bipartite disks replacing the marmoset portraits. The disks were black on the left and light grey on the right or vice versa. These two versions of the bipartite disks were presented randomly interleaved for 100 ms or 300 ms, two portrait exposure times that had prompted clear gaze following in the main experiment. Against the backdrop of the preceding considerations, we had hypothesized that the animals might prefer the target on the side of the darker half of the bipartite disk for both exposure times. However, contrary to our expectation, the animal preferred the target on the side of the brighter half of the disk, independently if positioned on the right or left side and, moreover, only for 100 ms exposure time. For 300 ms choices did not exhibit any preference (Figure 2B). This result does not support the hypothesis that marmoset gaze following is determined by a simple mechanism, confined to the comparison of the two ear tuft areas. It rather suggests that additional features such as the orientation dependent position and shape of the paler center parts of the face may matter as well.

### Congruent choices are accompanied by faster reaction times already at short exposure times

In the main experiment, the latencies of saccades indicating congruent choices were shorter than the ones for incongruent choices for exposure times up to 400 ms duration (Figure 3). Actually, this facilitation effect was strongest for the shortest exposure time, gradually decreased with exposure time and no longer reached significance for the longest durations tested (500 and 600 ms; see figure 3 legend for statistics), consistent with a time course of reflexive rather than volitional orienting. A similar facilitation effect for comparably short exposure durations has been seen in studies of macaque monkeys [11] and humans [8]. However, these studies did not report a gradual increase of reaction times with the time of exposure seen in our experiments on marmosets. This difference might be a consequence of the specific paradigm we used. In our experiments, the animals had to choose between two targets of equal appearance, rather than to follow the other’s gaze to a specific target as in the work on macaque monkeys and humans. Hence, our animals may have tended to extract additional information from the other’s face beyond gaze direction in an attempt to facilitate their choices, provided that this portrait was available long enough. This increased interest in the other’s face, gated by longer exposure times, can be expected to compromise the ability to quickly disengage attention at the time of the go-signal. This interpretation is supported by the experiments with control stimuli and the eye movements prompted by the appearance of the portraits we discuss below.

**Figure 3.**
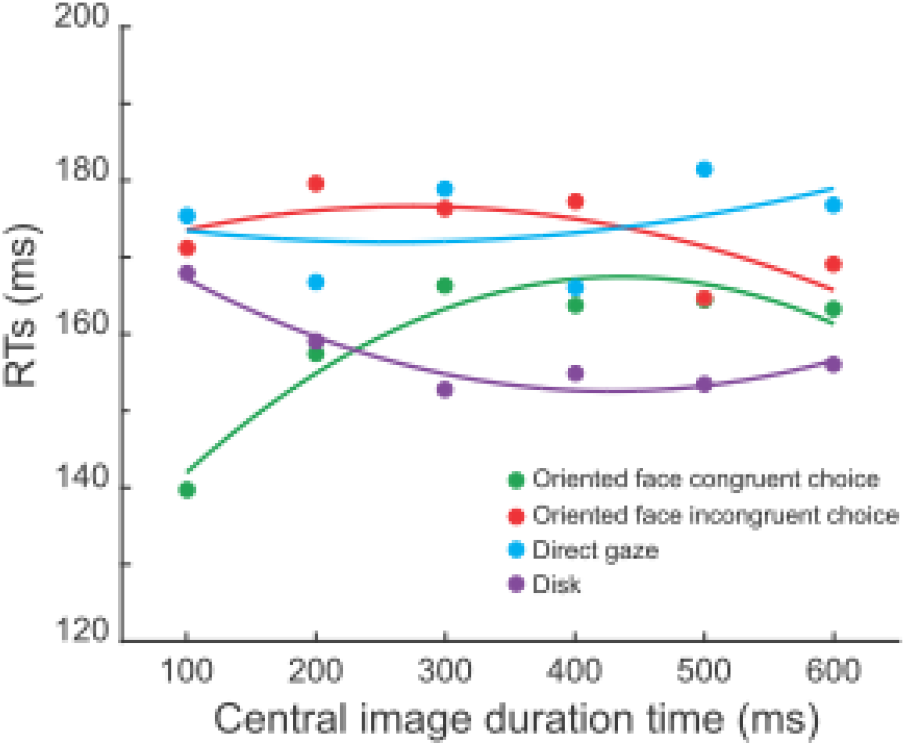
Oriented faces speed up reaction times for congruent choices. Plots of saccadic reaction times as function of the duration of presentation of the central stimuli. Data pooled over the three animals (monkeys M1, M2, M3) and choice direction. Saccadic reaction times indicating congruent choices were significantly shorter compared to the incongruent ones up to 400 ms of presentation duration (Wilcoxon rank-sum test, 100 ms: zval = −4.8221, p < 0.001; 200 ms: zval = −3.8449, p < 0.001; 300 ms: zval = −2.0341, p = 0.04; 400 ms: zval = −2.4745, p = 0.01). The individual plots are fitted with 2nd degree polynomial functions in an attempt to improve the visibility. The two fit that showed a significant dependence of saccade latency on the presentation duration of the central stimulus were the one for congruent choices prompted by oriented faces and the one for neural disk stimuli. The former exhibited a gradual increase with duration from a substantially shorted reaction time for a duration of 100ms (adjusted r^2^ = 0.86). The latter, on the other hand, a gradual decrease with duration (adjusted r^2^ = 0.89).

Saccades associated with the straight ahead face (“direct gaze”) exhibited latencies that were not different from the ones associated with incongruent choices to oriented faces. Interestingly, latencies of saccades associated with neutral disks showed an influence of exposure time that was qualitatively opposite to the influence on saccades for congruent choices: while being similar to saccades for straight faces for short presentation durations, they became shorter with increasing exposure time (see figure 3). The same held for the bicolor disk control stimuli (see figure S2). These results indicate that for marmosets, the attraction of the other’s face and not to non-biological stimuli increases with exposure time and correspondingly attentional disengagement is delayed.

Neutral objects were associated with relatively long saccadic reaction times when presented briefly, probably because of the need to scrutinize the object in order to assess its significance. Once its irrelevance is established after some 200 ms of presentation, the observer disengages his attention in order to prepare a fast saccadic choice. A short exposure to the oriented face can cause a profound shortening of saccadic reaction time, because the drive to follow gaze direction is already fully expressed whereas facial attraction is still building up. The idea that the development of facial attraction and in general the perception of faces may need much longer is also supported by a consideration of the pattern of saccadic exploration of the portraits (see supplementary figure S3 for details) whose complexity keeps growing with exposure time. Hence, the question is why the drive to follow gaze is fully expressed in saccadic reaction times for short exposure times, arguably too short to allow a detailed scrutiny of the face whereas the choice bias increases further with exposure time for up to 300 ms. We think that this dissociation between reaction times and choice probabilities might reflect the concerted action of two systems controlling gaze following. The first is fast, probably subcortical, controlling gaze following based on a rough and potentially error prone analysis of the other’s face, too limited to provide information on other aspects of the face like the identity or mood of the agent. With longer exposure and concomitantly processing time, this information becomes available, on the one hand binding attention but, on the other hand, also improving the directional precision of decisions.

### Concluding remarks

Gaze following is prevalent among numerous species but its strength and flexibility varies substantially between them [18]. As shown here, gaze following is also well developed in common marmosets, a new world monkey species. Marmoset gaze following is characterized by strong similarities with the gaze following behavior of the two old world primate species studied extensively, macaques and humans. The strongest argument for correspondence is the similar dependence on the time of exposure to the other’s gaze direction. In all three species the other’s gaze biases decisions on potential targets already after only 100 ms of exposure to the other’s gaze, too short to accommodate a more detailed scrutiny of the other’s face. However, given more time to explore the other’s face, the bias gets stronger in all three, in line with the assumption that primate gaze following is a faculty, consisting of an early reflex-line component that is complemented by a later, more flexible component, arguably also responsible for the more sophisticated emotional and cognitive control known to modulate gaze following [19–21]. The behavioral similarities between the gaze following behavior of marmosets, macaque monkeys and humans are in principle in line with the assumption of a homologous faculty, already available before the split of the new and old world monkey primate lines. This conclusion may strengthen the view that the marmoset may indeed become a useful model system for research into the underpinnings of disturbed human social interactions like autism, related to deficient gaze following and joint attention [22]. However, although compelling, the behavioral similarities established in our study may as well reflect behavioral convergence. Hence, comparative physiological and genetic studies of the underlying neural systems will be needed to strengthen the case for homology.

## Acknowledgments

This work was supported by a grant from the Deutsche Forschungsgemeinschaft (TH 425/12-2).

## SUPPLEMENTARY MATERIAL

### EXPERIMENTAL MODEL AND SUBJECT DETAILS

#### Common Marmosets

We trained 3 adult common marmoset monkeys (Callithrix jacchus; two females and one male, aged 7 years) to voluntarily enter a custom made monkey chair by means of positive reinforcement training and to accept the restriction of head movements through a head holder. Animals were all born in captivity and kept in a marmoset husbandry at approximately 26°C, 40%-60% relative humidity and a 12h:12h light-dark cycle. Access to water was always ad libitum, while food intake was controlled according to body weight (weight loss never less than 10% of the ad libitum weight) and amount of reward received in the experiment. Food consisted in fresh fruits and vegetables and standard commercial chow. As additional treats the animals received mealworms and locusts on days of good performance in the behavioral training and experiments. Reward given in the experimental setup consisted in self prepared marshmallow juice (1:2 marshmallows/water) with the addition of a small amount of gum arabic, or only gum arabic diluted in water, according to the individual animal’s preference. Experimental procedures were approved and supervised by the regional state authorities (Regierungspräsidium Tübingen and Landratsamt Tübingen, TVG N16/14) and are in agreement with the guidelines of the European Community for the care of laboratory animals.

#### Surgical and training procedure

All the animals underwent the surgical implantation of a titanium headpost under general anesthesia with sevofluran (2.5-5%), propofol (0.05-2 mg/kg/min), remifentanil (0.06-0.1 mcg/kg/min) and tight control of vital parameters. The headpost was fixed with three upside down T-shaped anchors, whose arms were placed between the skull and the dura. This was achieved by cutting a small slit into the bone using an ultrasound bone-knife (Mectron, Piezosurgery), allowing the insertion of the arms that were then rotated 90° under the bone. Two-component UV curing cement (ESPE Rely X Unicem 2) was used to close the bone slit and Super Bond C & B to glue the head post to the profiles protruding a few mm from the bone. After full recovery from surgery, the animals were gradually accustomed to head restraint through daily sessions of increasing duration, up to a maximum of 2 hours.

#### Experimental setup

The experiments were performed in a small sound proof room in daily sessions lasting between 30 min and 2 hours. The number of trials per session ranged from 50 (usually at the start of a session block after a few days break) up to 500 trials per session. The animals were sitting in a comfortable monkey chair that was placed on a table facing a computer screen (Beetronics, 10 Inch Monitor, 220 x 134 mm, 1920 x 1080 Hz resolution, framerate 60 Hz), at a distance of 32 cm. Eye movements were tracked with the EyeScan System ETL-200, through a camera placed on the right side of the screen, and resampled at 1 kHz. Reward was delivered by means of a small cannula placed in front of the animals’ mouth, on or very close to the upper lip, depending on the animals’ preference. The delivery of rewards was controlled via a pump, set to release one drop of fluid for each correct answer or more (2-3 drops), depending on the animal’s motivation.

#### Eye Position Calibration

Eye position was calibrated by asking the animal to pursue a human face (4×5°) that was slowly moving on a circular trajectory on the screen (circle diameter 5°) at a speed of 6 °/s. In order to prevent that the animal would lose interest in the face, we replaced it every 4 trials by another one, differing in identity and/ or expression. The animals followed a novel moving face spontaneously with smooth pursuit eye movements with interspersed catch up saccades allowing us to calibrate the eye position records by fitting the target trajectory to the eye trace.

#### Behavioral paradigm

Each trial started with the appearance of a small red dot (0.2°) in the center of the screen on a white background, available for a maximum of 500 ms to start fixation (fixation window (2° × 2°). Otherwise it disappeared and the trial was discarded. However, if fixation was acquired and maintained for 500 ms, the dot was replaced by the portrait of a conspecific portrait, in the main experiment randomly oriented towards a position at −5° or 5° on the horizontal, in 50% of the trials to the left side and in 50 % to the right side. In the control experiments the oriented faces were replaced by a face of a marmoset looking straight at the experimental animal, a monochromous disk (black or grey), or a bipartite-monochromous disk (left half black/right half grey or viceversa) respectively. Animals could freely explore the central images, as long as they kept the eye within the fixation window, which whose size corresponded to the image. Central images were presented for a variable duration of 100, 200, 300, 400, 500 or 600 ms. At the end of the image presentation time, the central image disappeared while at the same time a pair of 2 peripheral targets, human faces, looking straight and exhibiting a neutral expression (size 2 × 3°) appeared at +5° and −5° from the center. The identity of the animal presented in the center and the identity of the pairs of human faces serving as targets were kept constant. The appearance of the targets served as go signal, telling the animals to perform a saccade to one of the two targets. All choices were rewarded, as long as the indicative saccade landed within a window of 2° × 3°centered on the targets and was not carried out later than 500 ms after the go signal. Intertrial interval were kept constant at 1 second.

### STIMULI

The face stimuli used were based on photographs of the faces of marmoset conspecifics and humans that had been taken with a digital camera (Canon, Legria HFS30) and manually processed in Adobe Photoshop to unify their size and luminance. For the oriented face condition we used two different portraits of the same animal. The direct gaze stimuli were generated removing the peripheral white ear-tufts. The inner face feature were maintained and the resulting image was rescaled such as to match the spatial dimensions of the disk control stimuli.

### SACCADE IDENTIFICATION PROCEDURE

Saccades were identified by a Matlab routine as events characterized by an increase in instantaneous eye velocity above a threshold of 20 °/s. The performance of the algorithm was double checked by eye in order to discard false hits. As expected selected saccades respected the main sequence, i.e. relationship between amplitude and velocity / duration.

### STATISTICAL ANALYSIS

#### Binomial distributions of choice behavior and reaction times

For the statistical analysis of the binary decisions of the experiments animals, all sessions per animal and condition were pooled, yielding a binomial distribution allowing the detection of significant deviations from chance level (50%). The pairwise comparison of the binomial distributions for individual animals was based on chi-square tests which were carried without Yates correction, given that the number of trials per condition was large (>200). Pooled reactions times were compared between the various conditions by Wilcoxon-test with Bonferroni correction.

---

**Figure S1.**
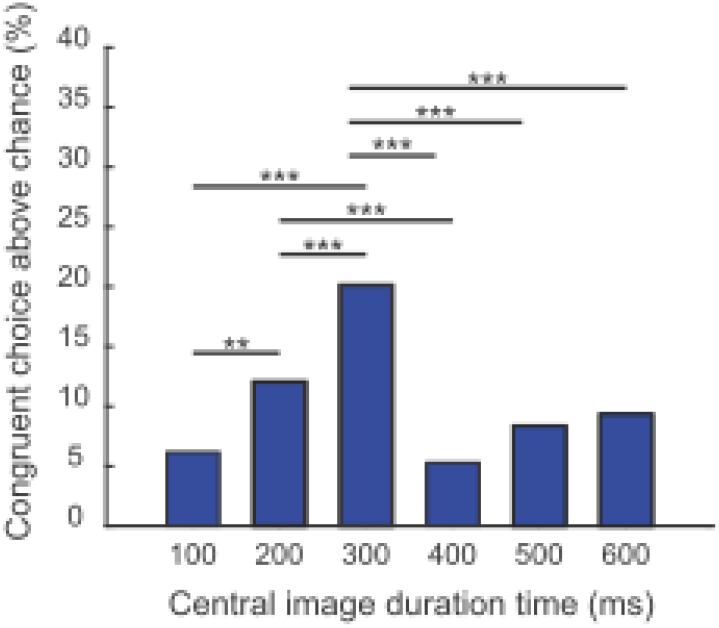
An exposure duration of 300 ms to the oriented face prompts the maximal gaze following. Bar chart of the number of congruent choices above chance level for the data shown in the left panel of figure 1B with statistical comparisons between presentation durations time view based on chi square tests without Yates correction, *** p < 0.001; ** p < 0.01; only significant comparisons shown). The percentage of congruent choices at 300 ms is significantly larger than for shorter or longer presentation durations.

**Figure S2.**
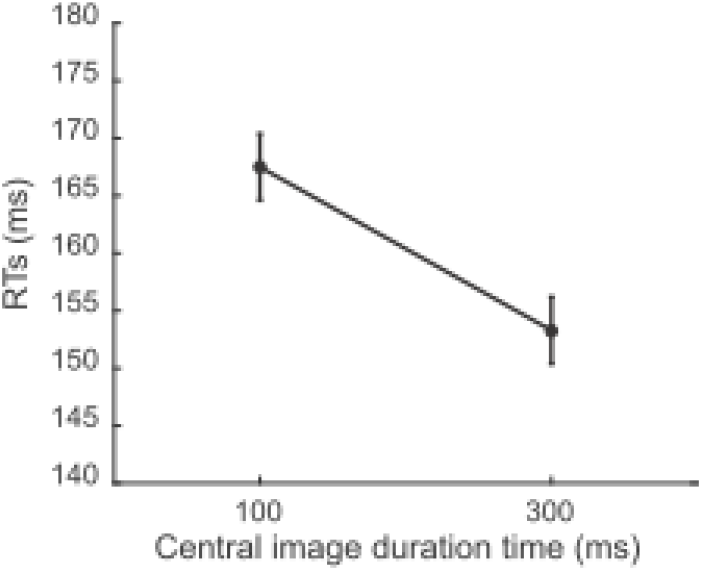
Saccadic reaction times for choices prompted by the bicolor disk control stimuli. No differences in saccadic reaction times (RT) were registered between choices towards the brighter and darker side (Wilcoxon rank-sum test, 100 ms: p = 0.8, zval = 0.235; 300 ms: p = 0.06, zval = −1.876). Hence, we pooled the both in order to assess the influence of presentation duration. As for the monochromous disk (see figure 3), RTs decreased with longer exposure to the stimulus (Wilcoxon rank-sum test, p < 0.001, zval = 3.776).

**Figure S3.**
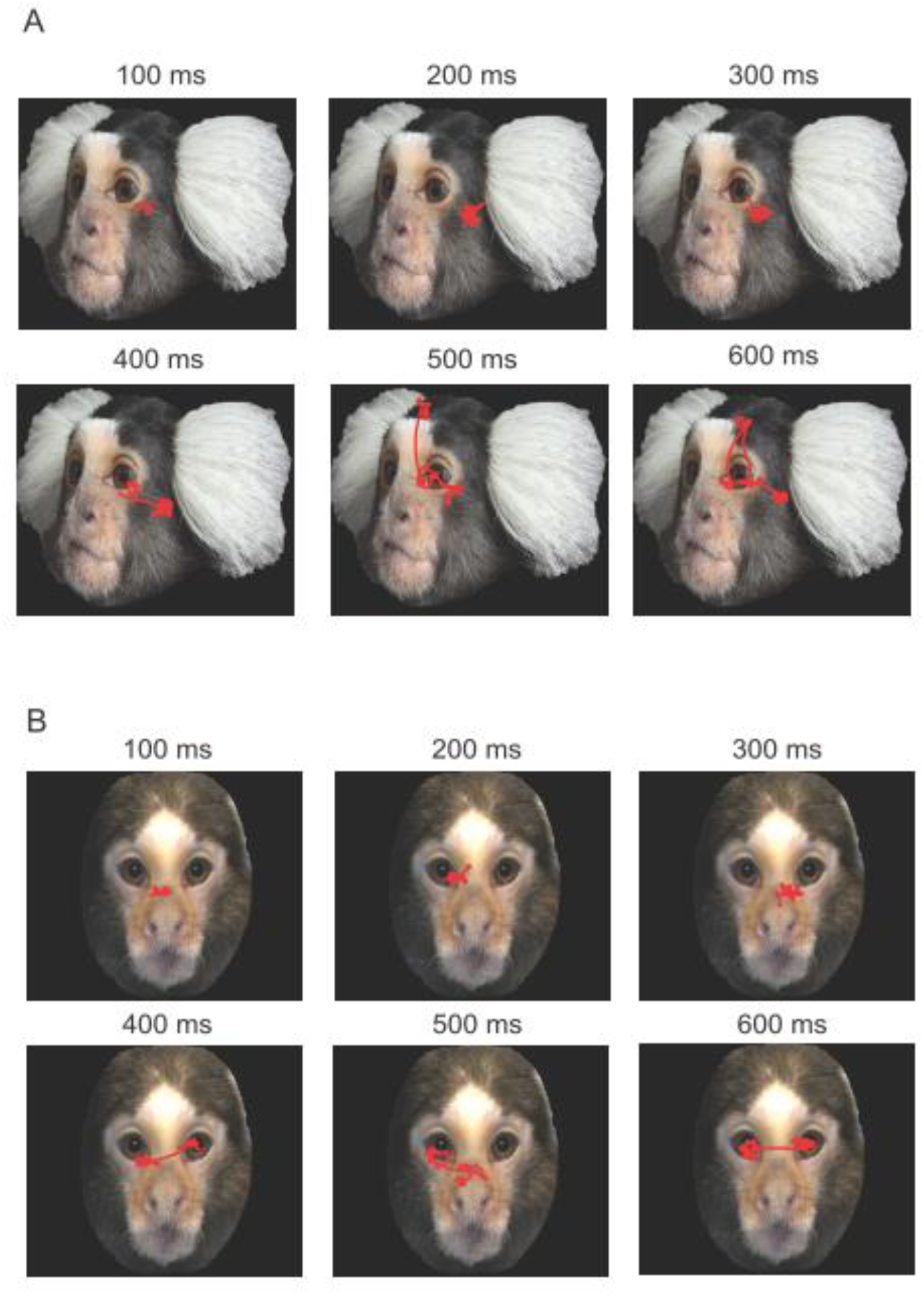
Only longer exposures to the other’s face allow the scrutiny of relevant facial features. Exemplary patterns of eye movement made by the observers when exposed to the oriented face of a conspecific (A) or alternatively to the frontal face of a conspecific lacking the white ear tufts (B) for different durations. Up to 300 ms the eyes of the observer stayed in a small, region of the face corresponding to the center of the image, arguably behaviorally not particularly relevant. Only exposure durations of 400 ms and longer allowed exploratory saccades, in these and most other cases aiming at the eye region and only rarely oriented towards the white ear-tuffs. The data are from individual sessions with monkey M2.

